# Nicotinamide restricts neural precursor proliferation to enhance catecholaminergic neuronal subtype differentiation from mouse embryonic stem cells

**DOI:** 10.1101/2020.05.13.094110

**Authors:** Síle M. Griffin, Mark R. Pickard, Clive P. Hawkins, Adrian C. Williams, Rosemary A. Fricker

## Abstract

Emerging evidence indicates that a strong relationship exists between brain regenerative therapies and nutrition. Early life nutrition plays an important role during embryonic brain development, and there are clear consequences to an imbalance in nutritional factors on both the production and survival of mature neuronal populations and the infant’s risk of diseases in later life. Our research and that of others suggest that vitamins play a fundamental role in the formation of neurons and their survival. There is a growing body of evidence that nicotinamide, the water-soluble amide form of vitamin B_3_, is implicated in the conversion of pluripotent stem cells to clinically relevant cells for regenerative therapies. This study investigated the ability of nicotinamide to promote the development of mature catecholaminergic neuronal populations (associated with Parkinson’s disease) from mouse embryonic stem cells, as well as investigating the underlying mechanisms of nicotinamide’s action.

Nicotinamide selectively enhanced the production of tyrosine hydroxylase-expressing neurons and serotonergic neurons from mouse embryonic stem cell cultures (*Sox1*GFP knock-in 46C cell line). A 5-Ethynyl-2’-deoxyuridine (EdU) assay ascertained that nicotinamide, when added in the initial phase, reduced cell proliferation. Nicotinamide drove tyrosine hydroxylase-expressing neuron differentiation as effectively as an established cocktail of signalling factors, reducing the proliferation of neural progenitors and accelerating neuronal maturation, neurite outgrowth and neurotransmitter expression.

These novel findings show that nicotinamide enhanced and enriched catecholaminergic differentiation and inhibited cell proliferation by directing cell cycle arrest in mouse embryonic stem cell cultures, thus driving a critical neural proliferation-to-differentiation switch from neural progenitors to neurons. Further research into the role of vitamin metabolites in embryogenesis will significantly advance cell-based regenerative medicine, and help realize their role as crucial developmental signalling molecules in brain development.

## Introduction

There is a wealth of evidence indicating that diet and nutrition play a key role during sensitive windows of brain development, when early organizational processes such as differentiation and maturation of specific neuronal pathways are underway (1-3). Each step of neuronal differentiation, including neural progenitor differentiation, neuronal fate specification, maturation and survival of developing neurons is highly regulated by extrinsic and intrinsic factors (4); with a lack of or an excess of nutritional elements leading to abnormalities in brain development (5-7).

Pluripotent stem cells provide an important *in vitro* model system to investigate early events during human development and the therapeutic use of stem cells is a promising approach to combat neurodegenerative processes in the brain, e.g. the replacement of midbrain dopamine neurons in Parkinson’s disease (PD) (8) or serotonergic neurons in neuropsychiatric disorders (9). However, successful exploitation of stem cell derivatives requires the ability to restrict stem cell proliferation linked to tumour formation, and to direct differentiation of stem cell candidates to higher and purer yields of desired cell phenotypes (10). The dopaminergic neurons of the nigro-striatal system that are affected in PD, and the serotonergic neurons that project to cortical regions and which are affected in neuropsychiatric disorders, develop in close proximity to the ventral midbrain (11). Therefore, early neurogenesis of these specific neuronal subtypes may be influenced by similar patterning signals. While a number of these signalling pathways have already been identified (e.g. Lmx1a (12), Pitx3 (13), Nurr (14)), it is likely that there are as yet undiscovered factors that modulate the fate of specific midbrain neuronal cell populations during development.

The developing brain is metabolically highly active, and changes in metabolism are known to influence neuronal development (15). Nicotinamide, the amide form of vitamin B_3_ (niacin), is a key molecule whose levels are tightly governed by cellular metabolism, and is a key factor in the metabolic pathway to produce nicotinamide adenine dinucleotide (NAD+), which is known to be essential for energy production in the cell (16). Optimal NAD levels are critical in preventing impaired neuronal metabolism due to mitochondrial dysfunction. An NAD-deficiency is a likely key-event in the pathogenesis of PD (6). Thus, restoring NAD levels through supplementation with precursors such as nicotinamide has the capacity to improve mitochondrial function, prevent NAD deficiency and promote neuroprotection and neuronal development in neuronal populations (5, 7, 17-19). In this context, nicotinamide has been used to promote differentiation of pluripotent cells under a wide variety of culture conditions (20-26). A previous study in our laboratory demonstrated the benefits of applying nicotinamide as a differentiation agent to aid the conversion of stem cells to mature GABAergic neurons (18). Findings from this work and published literature (27-29) imply that this bioactive nutrient may also function as a catecholaminergic differentiation signal implicated in the development or maintenance of basal ganglia circuitry.

Interestingly, a high level of nicotinamide obtained from a modern Western diet abundant in meat and vitamin supplements has been proposed as a nutritional factor that, in excess may predispose dopaminergic neurons to mitochondrial stress and subsequent neuronal apoptosis, leading to PD (5, 6). In support of this theory, excess nicotinamide administered postnatally to mice caused a reduction in dopamine levels (potentially through Sirt1 inhibition) (30). Furthermore, previous work in our group demonstrated that cultured stem cell–derived neurons respond positively to supplementation with nicotinamide within a dose range of 5 to 10 mM *in vitro* (18), however 20 mM dose induced cytotoxic effects on cells within 3 days of application (7), implying that levels of this vitamin require tight regulation in order to support and sustain normal neuronal functioning. Conversely, in human Pellagra, severe tryptophan/niacin deficiency leads to range of symptoms including dermatitis, diarrhoea, dementia, depression and other features of neurological disorders including Parkinsonism (16, 31). Alterations in nicotinamide levels have been linked with Alzheimer’s disease and Huntington’s disease (reviewed in (32)), and nicotinamide treatment in animal models has shown amelioration of neurodegeneration and associated behavioral recovery (33, 34).

The aim of the current study was to investigate whether nicotinamide, within a defined dose range, was able to influence the differentiation of embryonic stem cells into mature catecholaminergic neuron subtypes. Nicotinamide was applied to differentiating mouse embryonic stem cells (mESC; *Sox1*GFP knock-in 46C cell line (35)) during their conversion towards a neural fate. Cells were assessed for changes in their proliferation, differentiation and maturation; using immunocytochemistry and morphometric analysis methods. This study also focused on elucidating the mechanism(s) mediating neural specification by nicotinamide - that is, induction of cell-cycle exit and/or selective apoptosis in non-neural populations. Here we show that nicotinamide, at a specific dose and exposure time, caused an accelerated passage of pluripotent cells through lineage specification and further to non-dividing mature catecholaminergic neural phenotypes. This places nicotinamide as an affordable and effective signalling factor for efficiently deriving enriched catecholamine neurons, and marks this bioactive molecule as worthy of further investigation to clarify its role in normal brain development.

## Materials and Methods

All reagents used in this study were from Sigma, UK unless specified otherwise.

### Embryonic Stem Cell Culture

The mESC line 46C (a kind gift from Professor Meng Li, Cardiff University, UK) carrying a green fluorescent protein knock-in reporter targeted to the *Sox1* promoter (transiently expressed during the neural progenitor stage) was used throughout this study. mESCs were cultured in Glasgow Modified Eagles Medium (Invitrogen, UK) with the addition of 10% FCS, 0.1 M β-mercaptoethanol, 1 mM _L_-glutamine, 10 mM non-essential amino acids, 100 mM sodium pyruvate and 100 U/ml leukaemia inhibitory factor. Undifferentiated cells were routinely passaged every two days, re-plating at a density of 1 x 10^6^ cells/cm^2^. mESCs were maintained on 0.1% gelatin-coated tissue culture plastic at 37°C in a 5% CO_2_ incubator.

### Neural Differentiation

Monolayer neural differentiation was based on a previous protocol (13). For neural induction, undifferentiated mESCs were plated at a density of 9 x 10^4^ cells/cm^2^ in wells of a 0.1% gelatin-coated 6-well dish (day 0) in N2B27 serum-free medium: DMEM/F12 (Invitrogen), Neurobasal media (Invitrogen), 0.5% N2 (Fisher Scientific, Loughborough, UK), 1% B27 (Fisher Scientific), 2 mM _L_-glutamine and 0.1 μM β-mercaptoethanol. Medium was refreshed every other day. On day 7, 3 x 10^4^ cells were replated in 30 μl microdrops of N2B27 medium on 13 mm glass coverslips (Fisher Scientific) pre-treated with poly-L-lysine (PLL) (10 μg/ml) and laminin (2 μg/ml) in 24-well plates. After 4–6 h incubation, the wells were supplemented with N2B27 medium, refreshed every other day until day 14. Cells were treated with nicotinamide (10 mM), between days 0 and 7, and assays were performed on days 7 and 14. Control groups were not treated with nicotinamide.

### Viability Assay

A Countess™ automated cell counter was used to determine total cell counts and viability of mESCs. For a typical cell count, 10 μl cell suspensions was mixed with 10 μl 0.4% trypan blue stain, and 10 μl was added into a chamber. The image was focussed, and viable and non-viable cells were counted. Three counts per sample were averaged and cell viability was calculated as a percentage of the total number of cells.

### Apoptosis Assay

The Click-iT^®^ TUNEL Alexa Fluor^®^ imaging assay (Invitrogen, Paisley, UK) was employed to assay nuclear DNA fragmentation by catalytically integrating fluorescein-12-dUTP at 3’-OH DNA ends, using the enzyme Terminal Deoxynucleotidyl Transferase (TdT) to generate TdT-mediated dUTP Nick-End Labelling.

Monolayer adherent cells were fixed in 4% PFA for 15 min at 4°C then permeabilized with 0.2% Triton X-100 in PBS for 5 min. Cells were rinsed, then the TdT reaction buffer was added to each coverslip for 10 min at RT. The coverslips were incubated for 60 min at 37°C in 100 μl TdT reaction cocktail, avoiding exposure to light, then rinsed with 3% BSA. A Click-iT^®^ reaction cocktail (100 μl) was added to each coverslip for 30 min at RT, protected from light, then the cells were rinsed. To perform dual labelling, the cells were blocked and permeabilized with 0.02% Triton X-100 and 5% NGS for 1 h. Immunocytochemistry with fluorescent antibodies was then performed as detailed below. TUNEL was combined with examination of cell nuclei integrity, and assessments of cellular morphology were performed using fluorescence and phase contrast microscopy. Apoptotic cells were identified as TUNEL-positive and exhibiting a pyknotic nucleus.

### Cell Proliferation Assay

To determine the proliferative effect of nicotinamide, a ClickiT ^®^ EdU (5-ethynl-2’-deoxyuridine) cell proliferation assay (Invitrogen, Paisley, UK) was used, in accordance with the manufacturer’s instructions. Adherent monolayer cells were pulse labelled with EdU (10 μM) for 1 h prior to cell fixation. Standard PFA fixation (4% formaldehyde) and detergent permeabilization (0.5% Triton X-100) were then performed to facilitate access of the detection reagent to DNA. For EdU detection, a Click-iT^®^ reaction cocktail was prepared and applied to the cells for 30 min. The cells were rinsed with 3% bovine serum albumin (BSA) in PBS, followed by PBS. To investigate whether nicotinamide had elicited an effect on the proliferation of undifferentiated mESCs, or differentiated neurons, dual labelling of cultured cells with EdU and primary antibodies was undertaken. Co-labelling of EdU with native *Sox1*GFP-expression was used to identify progenitor cells that had undergone mitosis.

### Immunofluorescence Staining

Differentiated cells were fixed with 4% paraformaldehyde (PFA) for 20 min at 4°C. Fixed cells were washed three times with Tris buffered saline (TBS). Non-specific binding was blocked and the cells were permeabilized with 0.02% Triton X-100 and 5% normal goat serum (NGS) (PAA, The Cell Culture Company, Somerset, UK), for 1 h at room temperature (RT). Primary antibodies diluted in 1% NGS blocking buffer were added to the cultured cells overnight at 4°C (Oct4, 1:100, Santa Cruz; β-III-tubulin 1:500, Covance; TH, 1:1000, Chemicon; Serotonin, 1:500, Abcam). Negative primary controls consisted of cells treated with blocking buffer without the addition of primary antibodies. On the next day, three TBS washes were applied to the cells followed by incubation with Alexa Fluor secondary antibodies, (Cheshire Sciences, UK) diluted to 1:300 in 1% NGS blocking buffer, for 2 h at RT. Cultures were washed three times with TBS. Coverslips were mounted onto microscope slides using Vectashield hardset mounting medium containing 4’, 6-diamidino-2-phenylindole (DAPI) (Vector Labs, Peterborough, UK) to counterstain cell nuclei, before visualisation the following day.

### Cell Quantification

Cell samples were visualised using fluorescence microscopy (Nikon Eclipse 80i microscope; Nikon UK Limited, Kingston upon Thames, UK) and images acquired using a Hamamatsu ORCA camera (Hamamatsu Photonics UK Limited, Welwyn Garden City, UK). NIS-Elements imaging software, version BR 3.2 (Nikon UK Limited) was used to manually quantify positive antibody labelling in DAPI-stained cultures. Independent experiments were replicated three times and three to four coverslips per group were counted within an experiment. Specific cell populations were analysed by capturing six to eight random fields per coverslip. Clusters containing dense populations of neural progenitor and neuronal cells were excluded from data collection, since these cell networks were not countable.

### Cell Morphology Analysis

Neuronal cells were photographed from eight to ten random fields per coverslip from three independent experiments using a high-power objective lens (i.e. x 40 lens). Neuronal cells within cell clusters were omitted from morphometric analysis. Four morphological parameters were investigated: 1) measurement of neurotransmitter content within neuronal cell bodies (soma) using Fluorescence Intensity measures (see below); 2) number of primary neurite branches; 3) length of the longest neurite (μm), and 4) total neurite extent (μm).

Neuronal processes greater than two cell diameters in length were considered as true neurites and neurites of labelled cells were manually traced using the ImageJ plug-in NeuronJ (version 1.4.2; NIH). Neurite length was defined as the distance from the soma to the tip of the longest primary neurite and the combined lengths of all neurites per cell were designated as total neurite length.

### Fluorescence Intensity Measures

Fluorescence Intensity (FI) measures were obtained to determine levels of protein expression of specific neuronal populations derived from *Sox1*GFP mESCs, using ImageJ image analysis software (version 1.45s; NIH). Cell samples were captured at fixed exposure settings using a Hamamatsu ORCA camera with NIS Elements imaging software. Cells located within clusters were excluded from the FI analyses. FI values were evaluated by converting each colour image to grayscale and calibrating images using an optical density step tablet. FI readings were then corrected for the background fluorescence.

### Statistical Analysis

Statistical analysis was performed using GraphPad Prism version 5.00 (GraphPad Software Inc., La Jolla, California, USA). Data plotted on graphs is expressed as mean ± standard error of the mean (SEM). Unpaired two-tailed t-tests were used to compare data from cultures treated with 10mM nicotinamide versus control cultures. A level of *p*<0.05 was used as the limit for statistical significance.

## Results

### Nicotinamide promotes TH-expressing neurons and serotonin neuron differentiation from ESCs, in the absence of exogenous inductive molecules

The effects of nicotinamide on the *Sox1*GFP knock-in 46C mESC line were investigated. In all experiments, undifferentiated cells were treated with 10 mM nicotinamide between days 0 and 7, and the cells were further differentiated up to 14 days. Factor-free neural differentiation of *Sox1*GFP mESCs (without inductive signalling molecules or exogenous growth factors), produced low numbers of TH-expressing neurons per culture, as revealed by TH immunocytofluorescence. Nicotinamide treatment elicited a significant increase in the percentage of TH^+^ immunoreactive neurons within the whole cell population in comparison to control cultures (t = 6.1; p<0.001; 4.2 ± 0.5% in nicotinamide treated vs. 0.6 ± 0.3% in untreated conditions (Fig 1 A-C).

**Fig 1.**
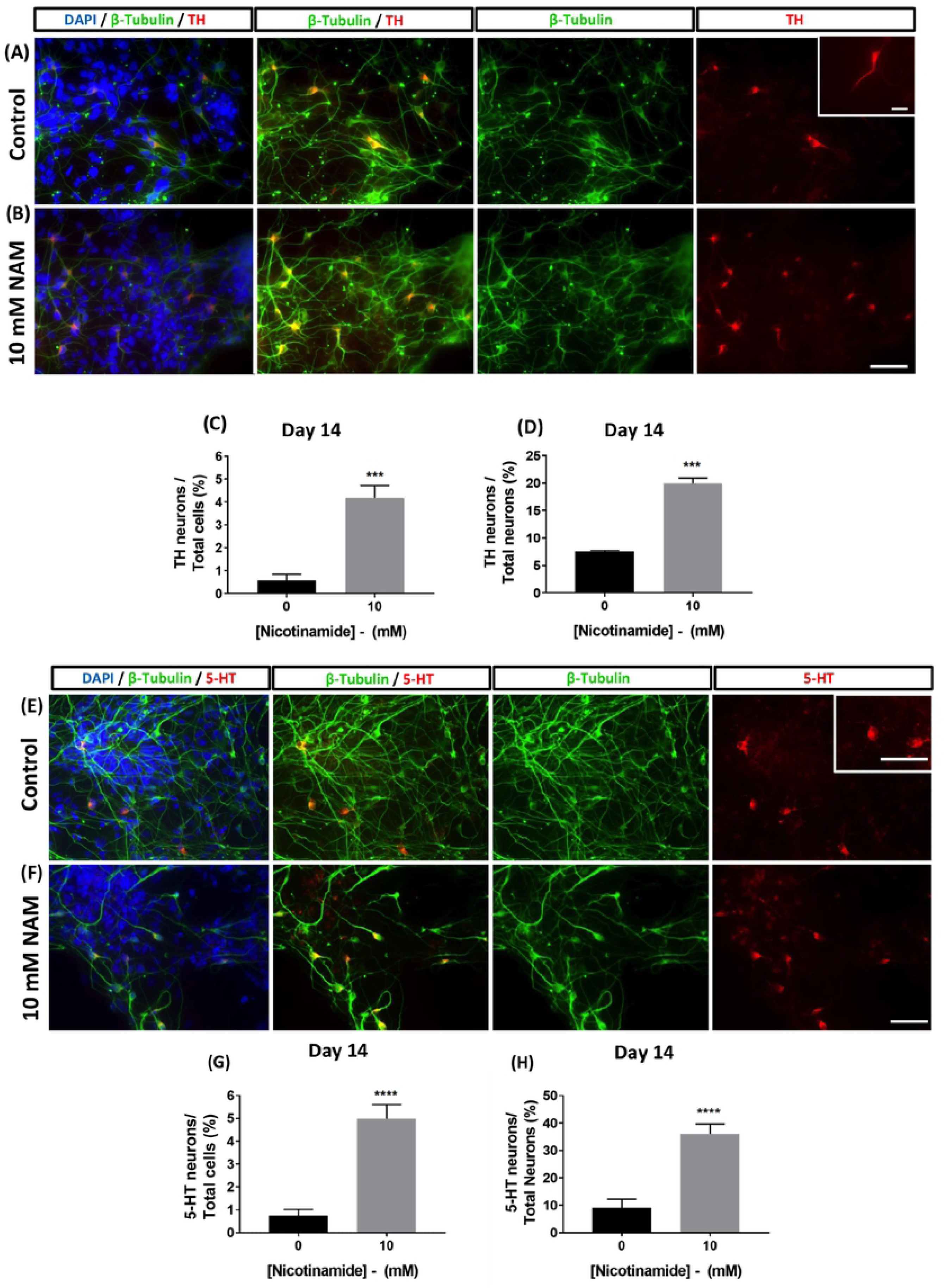
Nicotinamide promotes tyrosine hydroxylase-expressing neuron and serotonergic neuron differentiation from mESCs in the absence of exogenous inductive molecules. Nicotinamide added to cultures between days 0 and 7 significantly increased the percentage of tyrosine hydroxylase-expressing (A-D) neurons and serotonergic (E-H) neurons at day 14; both as a proportion of total cells (C,G) and as a proportion of βIII-tubulin-expressing neurons (D, H). Scale bar 50 μm applies to all images. ***p<0.001, ****p<0.0001

In addition, 10 mM nicotinamide treatment induced almost a three-fold increase in the percentage of neurons that were double-labelled for βIII-tubulin and TH (t = 13.89; p<0.001; 20.0 ± 0.9% in nicotinamide treated vs. 7.5 ± 0.2% in untreated conditions; (Fig 1D). In separate cultures, addition of 10 mM nicotinamide also induced an increased proportion of serotonergic neurons within the total DAPI+ cell population (t = 5.3; p<0.001; 5.0 ± 0.6% in nicotinamide treated vs. 0.7 ± 0.3% in untreated conditions; Fig 1 E-G). Cultures exposed to 10 mM nicotinamide contained a significantly greater proportion of neurons that were double-labelled for βIII-tubulin and 5-HT (t = 5.1; p<0.001; 36.1 ± 3.6% in nicotinamide treated vs. 9.2 ± 3.1% in untreated conditions; Fig 1H).

Taken together, these experiments indicate that in *in vitro* mESC cultures induced with a period of early exposure to nicotinamide there was both an enhanced yield and an enrichment of TH-expressing and 5-HT-expressing neurons.

### Nicotinamide increases the maturation and complexity of TH-expressing neurons

Next, the effect of nicotinamide on neuron development and maturation was investigated in mESC-derived TH-expressing neuron populations. Image analyses indicated that for TH-expressing cells, the average primary neurite length was notably longer in 10 mM nicotinamide conditions than controls (Fig 2 A,B).

**Fig 2.**
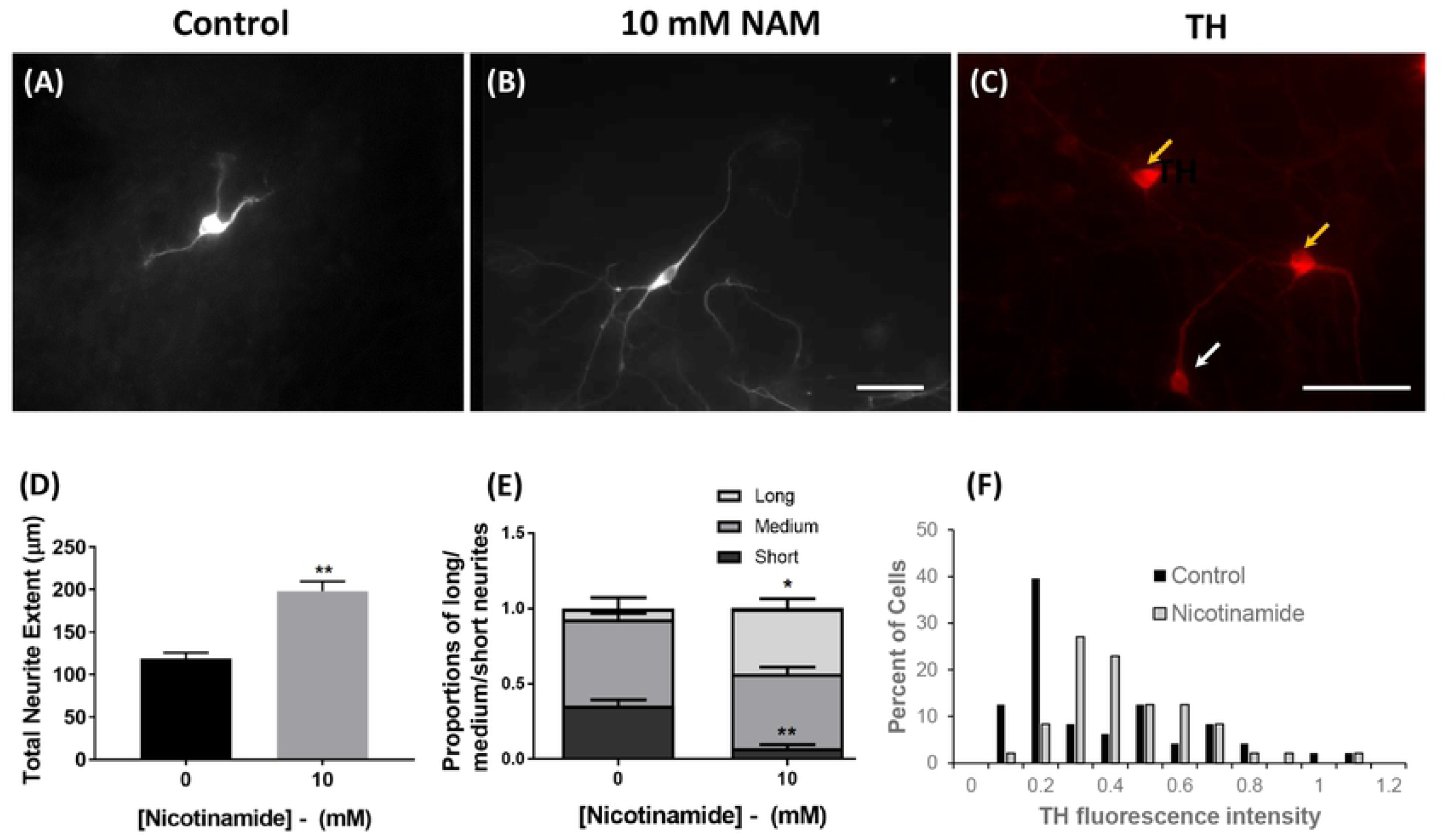
Nicotinamide enhances neuronal maturation in tyrosine hydroxylase-expressing neurons populations. The proportion of 46C-derived cells displaying “short” primary processes was significantly decreased in cultures exposed to 10 mM nicotinamide between days 0 and 7, concomitant with an obvious increased trend towards “longer” neurite processes, compared with controls (A-B, D-E). Within cultures, cells showed intense TH immunoreactivity (yellow arrows) or less strong TH expression (white arrow) (C). Addition of nicotinamide enhanced the number of neurons expressing strong TH immunofluorescence (F). Scale bars = 50 μm. **p<0.01, *p<0.05

Cultures treated with 10 mM nicotinamide also contained TH-expressing neurons with a significantly increased total length of all neurites (t = 5.9; p<0.01; 197.9 μm ± 11.6% in nicotinamide treated vs. 118.8 μm ± 7.1% in untreated conditions; Fig 2D). The numbers of neurites per neuron were not changed (t = 0.3; n.s; 2.0 ± 0.1 in nicotinamide treated vs. 1.9 ± 0.1 in untreated conditions).

Grouping the data into the proportion of TH-expressing neurons with short (≤ 60 μm), medium (60-120 μm) and long (≥ 120 μm) primary neurite processes, confirmed that the proportion of 46C-derived TH-expressing cells displaying “long” primary neurite outgrowths were significantly increased in cultures exposed to 10 mM nicotinamide (t = 4.0; *p<*0.05), concomitant with a significant decrease in the proportion of “shorter” TH^+^ primary neurites (t = 6.5; *p<*0.01; Fig 2E).

In addition, at 14 DIV there was a substantial increase in the number of TH-expressing cells displaying higher levels of TH fluorescence in nicotinamide-treated cultures compared with control conditions (Fig 2C). A histogram plot of TH immunofluorescence intensity indicated a shift from 0.1-0.2 units of intensity in the majority of TH-expressing neurons cultured in control conditions, to 0.3-0.6 units of intensity in the majority of TH-expressing neurons from nicotinamide treated cultures (Fig 2F), suggestive of increased levels of TH protein in the nicotinamide-treated cultures.

Therefore, following exposure to nicotinamide between days 0 and 7, mESC-derived TH-expressing neurons appeared to be more mature at day 14; they possessed longer primary neurites, they had an increased total length of neurites, and increased levels of the TH protein.

### Early nicotinamide treatment reduces total live cell numbers, but does not induce apoptosis or alter cell viability in monolayer cultures

Administration of 10 mM nicotinamide from day 0 to 7 of differentiation caused a significant reduction in the total number of cells by day 7 (Fig 3A,B,D), when compared to control conditions (t = 3.6; *p<*0.05; 4.33×10^5^ ± 3.50×10^5^ cells in nicotinamide treated vs. 2.83×10^6^ ± 4.77×10^5^, in control conditions). However, nicotinamide did not affect culture viability at this time-point (t = 1.3; n.s.; 81.5 ± 0.5% live cells in nicotinamide treated vs. 87.0 ± 3.2% in control conditions (Fig 3F).

**Fig 3.**
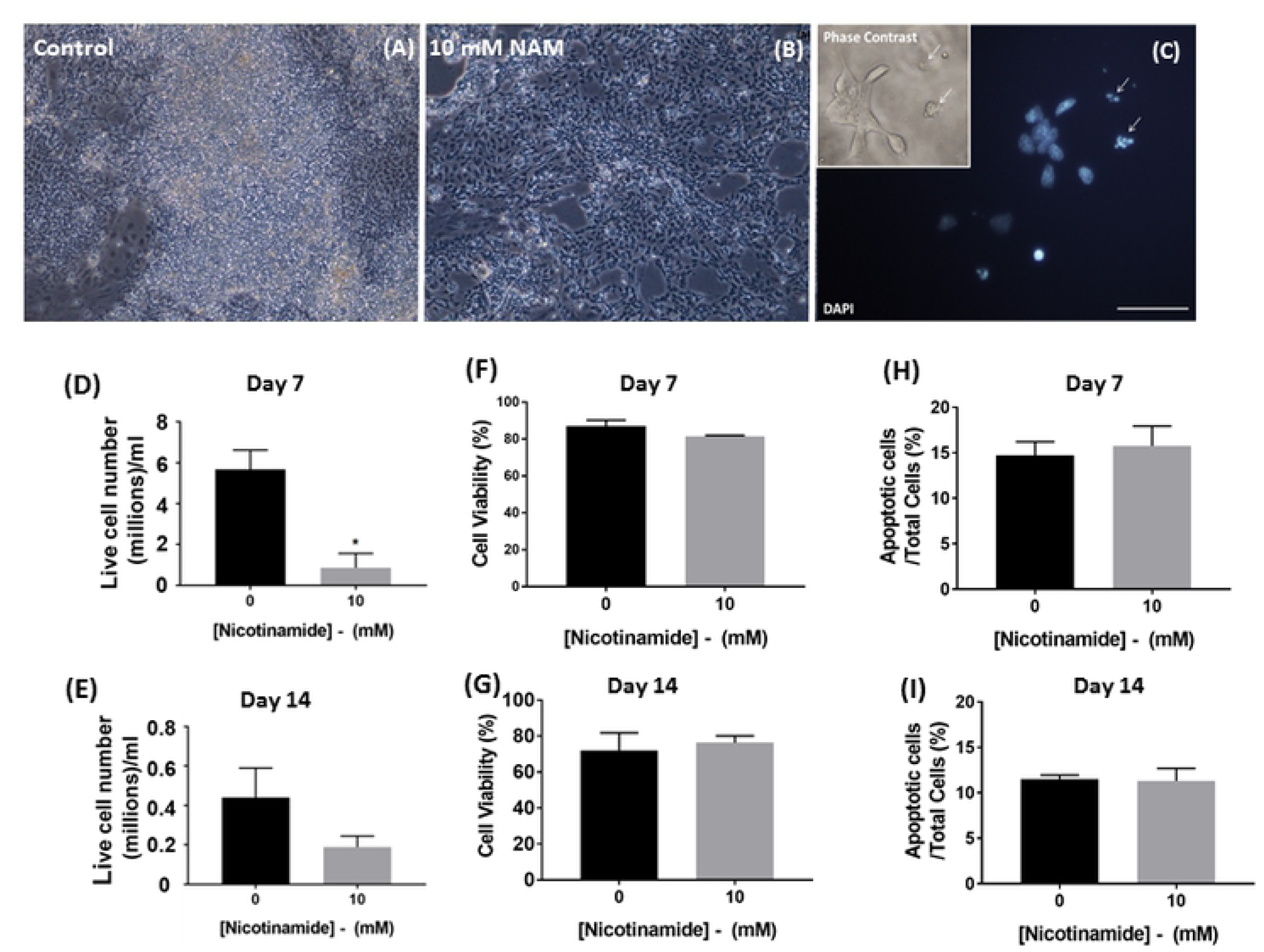
Nicotinamide does not induce apoptosis or alter cell viability in monolayer cultures. Monolayer cultures were exposed to nicotinamide between days 0 and 7, and viability assessed in terms of both total and percentage viable cells at days 7 and 14 of differentiation (A-G). Nicotinamide did not elicit any significant differences in the percentage of viable cells remaining in the cultures at days 7 and 14 (D-E). Nicotinamide administered during the first 7 days of differentiation induced a reduction in live cell numbers at 14 DIV, without evidence of toxicity (F-G). The percentages of apoptotic cell populations were not significantly altered by early nicotinamide treatment in adherent monolayer cultures on days 7 and 14, respectively. Scale bars = 50 μm. *p<0.05

These parameters were also addressed in cultures treated with nicotinamide from day 0-7 that were further differentiated up to a total of 14 days. There was a decrease in absolute cell numbers in the nicotinamide-treated cultures (t = 1.6; n.s.; 9.44×10^4^ ± 2.80×10^4^ in nicotinamide treated vs. 2.20×10^5^± 7.52×10^4^ in control conditions; Fig 3E), in agreement with previous findings (8). Similar to the situation on day 7, early nicotinamide administration did not alter the viability of cells by day 14 (t = 0.4; n.s.; 76.3 ± 3.8% viable cells in nicotinamide treated vs. 72.0 ± 9.8% in control conditions; Fig 3G).

The effect of nicotinamide on cell survival/death was further investigated using a TUNEL apoptosis assay, in combination with a morphological analysis of pyknotic cell nuclei (condensed or fragmenting morphologies) within the total cell population per microscopic field, at 7 and 14 days respectively (Fig 3C).

Nicotinamide addition during the early stages of differentiation did not significantly alter the percentage of apoptotic cells present on day 7, relative to untreated groups (t = 0.5, n.s.; 15.8 ± 2.1% in nicotinamide treated vs. 14.4 ± 1.4% in control conditions; Figure 3H). Similarly, on day 14, the percentage of nuclei exhibiting pyknosis and TUNEL staining did not significantly differ from control cultures (t = 0.1, n.s.; 11.3 ± 1.4% in nicotinamide treated vs. 11.5 ± 0.4% in control conditions; Figure 3I).

### Early nicotinamide administration halts proliferation of *Sox1GFP*-expressing neural progenitor cells

Due to differences in total cell number, the percentage of proliferating cells in cultures treated with nicotinamide from day 0-7 was compared with untreated controls. Cultures received 5-ethynyl-2’-deoxyuridine (EdU) for 1 h prior to cell fixation at the end of the 7 day culture period to determine the numbers of actively dividing cells after partial differentiation. This pulse-labelling experiment revealed that 61% of cells in cultures that had been differentiated without nicotinamide showed positive EdU uptake, whereas only 29% of cells in the cultures containing nicotinamide were labelled with EdU on day 7 (t = 10.6, *p<*0.001; 28.7 ± 2.9% in nicotinamide treated vs. 60.7 ± 0.9% in control conditions; Fig 4 A,B).

**Fig 4.**
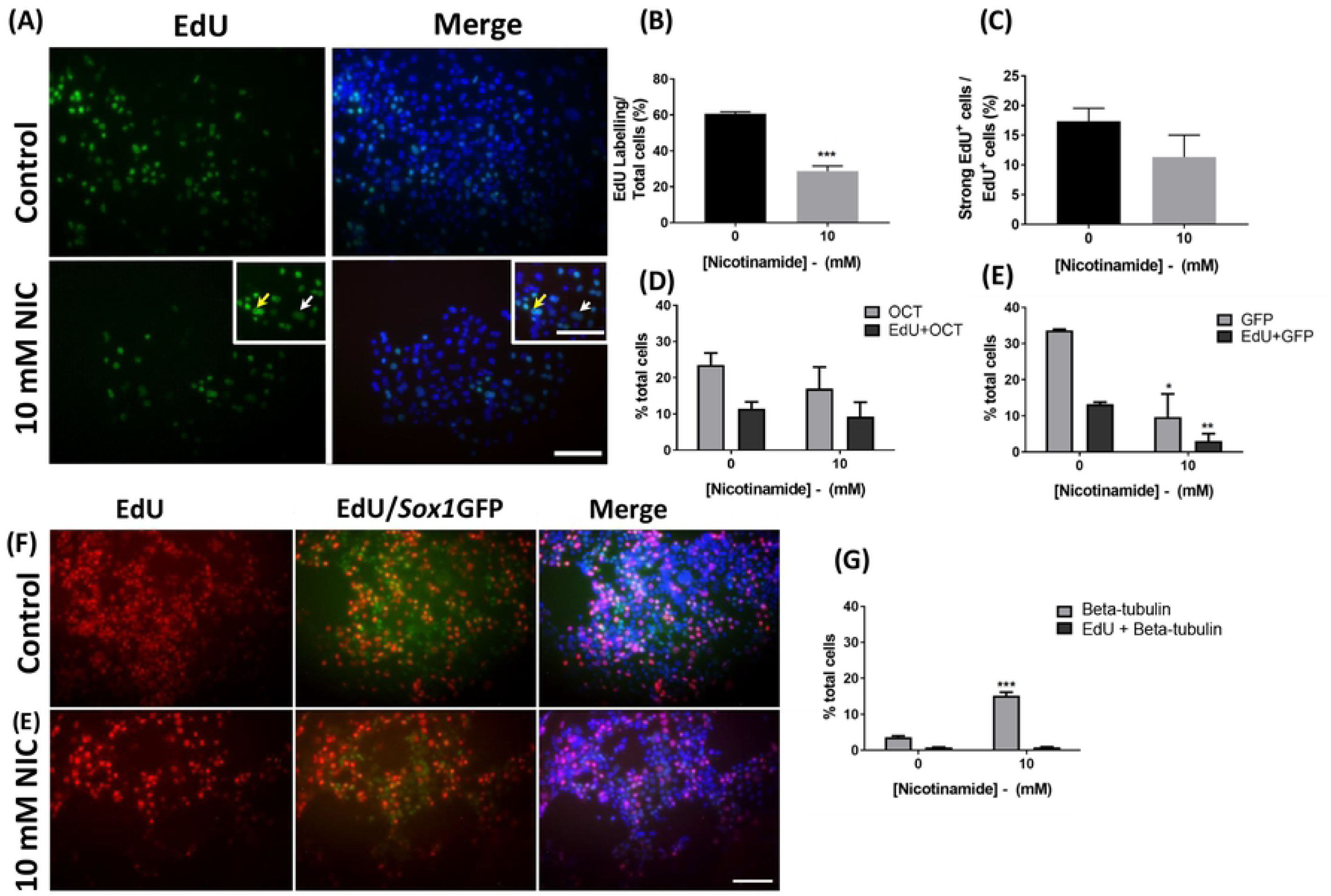
Early nicotinamide administration halts proliferation of *Sox1*GFP derived neural progenitor cells. A reduction in cell proliferation was observed in 10 mM nicotinamide-treated cultures versus control groups (A-B). There was no significant difference in the percentage of strongly labelled EdU^+^ cells in cultures treated with nicotinamide from day 0-7, in comparison to untreated groups (C). Scale bar = 50 μm for main images and 60 μm for inserts. Arrows indicate cells expressing strong EdU labelling (yellow) and weaker EdU labelling (white). Nicotinamide treatment caused a slight reduction in the percent of Oct4-expressing cells by day 7, but did not alter the percent of EdU-labelled cells within this population (D). Nicotinamide treatment significantly reduced the percentage of GFP-expressing cells, concomitant with a decrease in the percent of EdU-labelled cells in the *Sox1*GFP-positive population (E, F). Nicotinamide treatment significantly enhanced neuron-specific βIII-tubulin-expressing cells, without altering the percentage of EdU-labelled cells within the neuronal population (G). Scale bar 100 μm applies to all low magnification images. ***p<0.001, **p<0.01, *p<0.05 - comparing 10 mM nicotinamide treatment to the equivalent control conditions.

Nicotinamide has been previously found to accelerate neural fate commitment and in turn, neuronal maturation during early development (18). Therefore, the second aim of the proliferation study was to determine whether nicotinamide was directing cells to exit the cell cycle at an earlier time point than in untreated cultures. For this reason, we classified EdU-labelled cells as having either strong or weak incorporation of EdU in their nuclei. NIS-image software was used to characterize the labelled cells as exhibiting expression levels predominantly within either the intensity range of 240-260 units (“strong expression”), or at levels of 140-160 units (“weak expression”). The percentage of EdU-labelled cells in nicotinamide-treated cultures that exhibited “strong” expression levels was slightly reduced, but did not significantly differ from control cultures (t = 1.4, n.s.; 11.3 ± 3.6% in nicotinamide treated cultures vs. 17.4 ± 2.2% in control conditions; Fig 4C).

To gain further insights into the effects of nicotinamide on the cell cycle, the uptake of EdU was measured within specific cell populations (Oct4, *Sox1*GFP or βIII-tubulin-expressing cells) in control and treatment cultures. Addition of 10 mM nicotinamide between days 0 and 7 had no effect on the proportion of Oct4+ cells, (t = 1.0, n.s.; 16.9 ± 5.9% in nicotinamide treated cultures vs. 23.6 ± 3.3% in control conditions; Fig 4D); or the percent of EdU labelled cells within the Oct4+ population (t = 0.5, n.s.; 9.2 ± 4.0% in nicotinamide treated cultures vs. 11.5 ± 1.8% in control conditions).

In contrast, proliferation was significantly reduced in neural precursor cells after treatment with nicotinamide. The percentage of GFP+ cells was significantly decreased by day 7 (t = 3.8; p<0.05; 9.7 ± 6.4% in nicotinamide treated cultures vs. 33.6 ± 0.3% in control conditions; Fig 4 E,F). Similarly, the proportion of precursor cells co-localising *Sox1*GFP+ expression and EdU-labelling was significantly lower following nicotinamide treatment (t = 4.8, p<0.01; 3.0 ± 2.0% in nicotinamide treated cultures vs. 13.2 ± 0.6% in control conditions).

There was no detectable difference in the proportion of cells that co-localised EdU and βIII-tubulin between treated and untreated cultures (t = 0.2, n.s.; 0.8 ± 0.1% in nicotinamide treated cultures vs. 0.9 ± 0.1% in control conditions; Fig 4G).

### Nicotinamide and exogenous inductive factors act synergistically to enhance TH-expressing neuron differentiation from *Sox1*GFP mESCs

Nicotinamide treatment was then applied to a more advanced stem cell differentiation protocol with addition of the ventralizing factors Shh and FGF8 (36). Mouse ESCs were differentiated for up to a total of 14 days in a monolayer culture. Early supplementation with 10 mM nicotinamide (day 0-7) in addition to exogenous factors led to a significant increase in the percentage of TH-expressing immunoreactive neurons within the whole cell population (t = 11.1; *p<*0.001; 12.7 ± 0.4% in nicotinamide treated vs. 6.2 ± 0.4% in control cultures with exogenous factors but no nicotinamide; Fig 5). Under these culture conditions, individual βIII-tubulin expressing neurons were not quantified manually due to the high density of βIII-tubulin^+^ neuronal networks, making it impossible to quantify the number of TH+ cells as a proportion of total neurons. Nor was it possible to determine if there had been a specific enrichment of TH-expressing neurons within the total neuron population.

**Fig 5.**
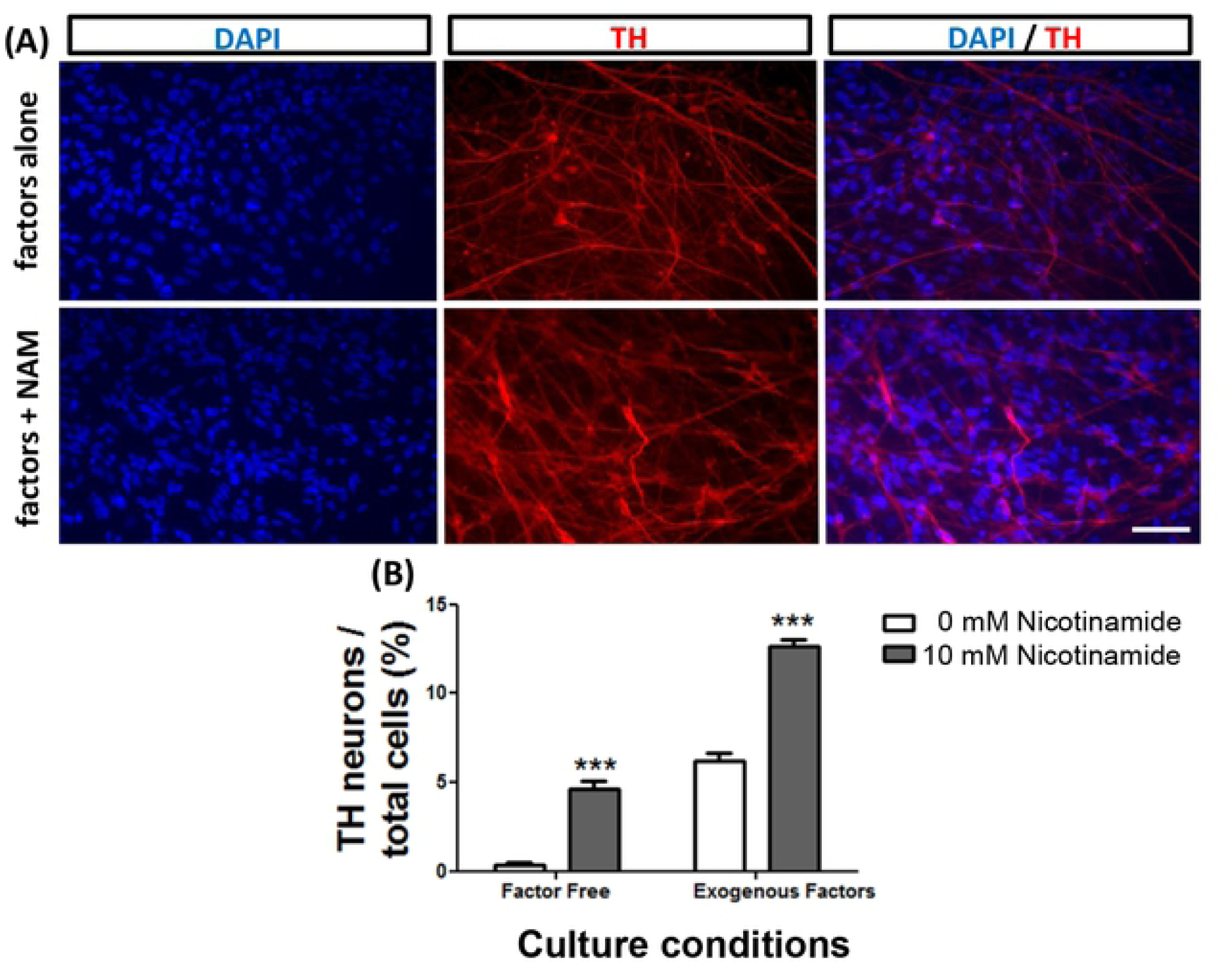
Nicotinamide and exogenous inductive factors act synergistically to enhance the production of TH-expressing cells from mESCs. The addition of exogenous factors to differentiating mESCs enhanced the production of dopamine neurons (A,B - white bars). Supplementation with nicotinamide induced similar TH neuron numbers as supplementation with exogenous factors; and both treatments together produced an additive effect to further increase TH neuron numbers (B - grey bars). Scale bar = 50 μm for all images. ***p<0.001-comparing 10 mM nicotinamide to 0 mM conditions.

## Discussion

The developing brain is influenced by genetic and environmental factors. There is a rapidly growing body of evidence that bioactive external signalling molecules such as vitamins exert a strong influence on neural development (18, 37); and an imbalance in their supply may contribute to a range of neurodevelopmental or neurological disorders later in life (32, 38).

Our previous work showed that nicotinamide could act as a morphogen to both enhance and accelerate the neural specification of mouse embryonic stem cells into neural progenitors, and subsequently as neurons (7, 18). The addition of nicotinamide to differentiating stem cells significantly elevated the numbers of neurons and GABAergic neurons generated *in vitro*, although the proportion of total neurons that were GABAergic was similar for both nicotinamide-treated and non-treated cultures, suggesting that nicotinamide was not acting selectively to enhance numbers of GABAergic neurons (18). This current study aimed to investigate whether nicotinamide might play a role in the specification of catecholamine neurons from mouse embryonic stem cells *in vitro*. In addition, the mechanism of action of nicotinamide in driving the neural and neuronal specification of stem cells was explored.

The results presented here reveal that 10 mM nicotinamide acted as a powerful morphogen to induce the differentiation of both TH-expressing and serotonin-expressing neurons from mouse embryonic stem cells. Nicotinamide’s mechanism of action was mediated through cell cycle exit rather than the selective apoptosis of non-neuronal cells. In addition, nicotinamide alone was sufficient to increase numbers of TH-expressing neurons from mouse ESCs to similar levels to those obtained using more complex signalling cascades, making this molecule an attractive, simple and cheap alternative for stem cell differentiation protocols.

The ESC monolayer differentiation method was chosen for this study as catecholaminergic neurons can readily be derived from *Sox1*GFP+ populations using this method (35), making it an ideal system with which to study the effects of nicotinamide. Furthermore, the monolayer cultures facilitated direct visualisation of the morphological changes of the ESCs during the course of their differentiation, allowing for the robust identification of any effects elicited by nicotinamide.

Addition of nicotinamide from days 0-7 in the 14-day differentiation protocol induced a significantly higher proportion of neurons expressing either TH or 5-HT. Thus, under serum-free and factor-free differentiation conditions, the enhanced differentiation of both TH-expressing and 5-HT-expressing neurons was due specifically to nicotinamide’s action on differentiating stem cells. Critically, unlike in our previous work that showed no specific enhancement of GABAergic neuronal differentiation (18); in the present study, nicotinamide enriched the proportion of total neurons that were either TH-positive or 5-HT-positive, i.e. this effect was subtype-specific. The fate of dopaminergic neurons has previously been shown to occur early in the conversion of ESCs into neurons, at or before the expression of *Sox1* in neural progenitors (11). This could explain why nicotinamide caused a selective enhancement of TH and 5-HT-expressing neurons in differentiated cultures, by influencing the fate of progenitors during early neural conversion.

Both TH and 5-HT-expressing neurons represent neuronal populations generated by similar ventralization signals originating from around the boundary of the midbrain and hindbrain (39). Therefore, the similar effects seen with *in vitro* differentiation of TH and 5-HT-expressing neurons mimic the differentiation pathways occurring during normal development *in vivo.* Nicotinamide appears to be important in directing both TH and 5-HT expression in neurons, suggesting that it is acting before the period when specific neuronal fates are being determined. Future work should investigate the combination of nicotinamide with Shh, FGF8 and noggin (a known agonist of bone morphogenic protein (BMP) and important for development of serotonergic neurons), which may lead to increased yields of serotonergic neurons derived from stem cells *in vitro*. Indeed, transplants of both dopamine and serotonin neurons have been advocated for individuals with Parkinson’s to ameliorate the motor and the non-motor symptoms as recent evidence indicates that the numbers of serotonin neurons decline in PD patients despite their receiving dopamine neuron grafts (40).

Importantly, the percentage of TH^+^ neurons generated in nicotinamide-treated cultures in this study (20%) approached the number shown previously to be generated in the presence of the ventralizing signals: sonic hedgehog (Shh) and fibroblast growth factor 8 (FGF8) (∼28%) (11, 41). This suggests that nicotinamide may have a similar potency to these well-established inductive signalling factors, thereby offering a cheap and simple alternative in differentiation protocols. Further, in the current study, a combination of all three factors: nicotinamide, Shh and FGF8, had an additive effect, more than doubling the yield of TH^+^ neurons. It is reasonable to conclude that nicotinamide may function synergistically with protein signalling molecules such as Shh and FGF8 during neural specification, to direct differentiating neural progenitors to adopt a dopamine cell fate, thus advocating the use of nicotinamide as a supplementary factor for current stem cell differentiation protocols.

The results presented here indicate that treatment with nicotinamide early in the differentiation of embryonic stem cells promoted neurite elongation in TH^+^ cells and the intensity of TH expression is in line with previous research examining other neuronal subtypes (18), thus suggesting an important role for nicotinamide in the maturation of TH-expressing neurons. We also observed an increased expression of the TH fluorescence in neurons derived from nicotinamide-treated mESCs. This elevated TH expression could imply that there is increased metabolic activity of the enzyme responsible for the production of the neurotransmitter dopamine, suggestive of increased maturation of the TH-positive neurons. In this regard, NADH is integral to the production of tetrahydrobiopterin, a co-factor necessary for tyrosine hydroxylase (29), the rate-limiting enzyme in catecholamine biosynthesis, also deficient in PD. Similarly, nicotinamide has been shown to elevate the expression of neurotransmitter within mESC-derived GABAergic neurons (18).

Consistent with our previously published data (7), nicotinamide did not affect cell viability at 7 and 14 DIV. Data from the current study suggest that whilst the N2B27 medium supports a level of continued proliferation of NPCs in cultures at day 7 of differentiation, exposure to nicotinamide during days 0-7 promotes the exit of NPCs from the cell cycle. Similar to the role of the vitamin A derivative, retinoic acid in neural development (42), this points to an important role for nicotinamide in the suppression of *Sox1*-positive precursor proliferation, promoting the transition from neural progenitors to neurons.

The results from the TUNEL assay indicated that the reduction in cell number in the presence of nicotinamide did not result from cell death in the cultures. This study investigated the action of nicotinamide within a dose range of 5-10 mM, previously shown not to cause toxicity in differentiating stem cell cultures (7). In support of this bioactive molecule acting to protect cells from death, nicotinamide is a naturally occurring inhibitor of the enzyme poly(ADP-ribose) polymerase-1 (PARP1) and application of nicotinamide has been shown to increase the efficiency of neuralization of human embryonic stem cells by rescuing neural progenitors from parthanatic cell death concomitant with an observed reduction in PARP1 activity (43). Conversely, EdU incorporation experiments indicated that a significantly lower number of *Sox1* positive NPCs were dividing on day 7 in the presence of nicotinamide. EdU expression indicated that labeled cells had exited the cell cycle immediately after incorporating EdU in S-phase. Since the vast majority of βIII-tubulin-expressing neurons were rarely double-labelled for EdU, it is reasonable to conclude that many of the differentiated neurons were already postmitotic at day 7 when EdU was applied to the nicotinamide-treated cultures.

## Conclusions

This study demonstrates that the small bioactive vitamin metabolite nicotinamide significantly enhances the differentiation of stem cells into TH- and 5-HT-expressing neurons. Nicotinamide’s strong influence on the development of the specific neuronal subtypes of catecholamine neurons suggests that it plays a critical role in phenotype-specific neuronal differentiation. It would be important to determine whether and how this vitamin derivative is an essential dietary component for normal brain development. The positive influence of nicotinamide in both the acceleration of differentiation and neuronal maturation is critically important, in the drive to increase the efficient production of subtype-specific neurons for the treatment of neurodegenerative conditions such as Parkinson’s disease as well as in neuropsychiatric disorders.

## Acknowledgements

We wish to thank Professor Meng Li, Cardiff, for the generous gift of the 46C embryonic stem cell line used in this study.

